# Characteristics of functional enrichment and gene expression level of human putative transcriptional target genes

**DOI:** 10.1101/085654

**Authors:** Naoki Osato

## Abstract

**Background:** Transcriptional target genes show functional enrichment of genes. However, how many and how significantly transcriptional target genes include functional enrichments are still unclear. To address these issues, I predicted human transcriptional target genes using open chromatin regions, ChIP-seq data and DNA binding sequences of transcription factors in databases, and examined functional enrichment and gene expression level of putative transcriptional target genes.

**Results:** Gene Ontology annotations showed four times larger numbers of functional enrichments in putative transcriptional target genes than gene expression information alone, independent of transcriptional target genes. To compare the number of functional enrichments of putative transcriptional target genes between cells or search conditions, I normalized the number of functional enrichment by calculating its ratios in the total number of transcriptional target genes. With this analysis, native putative transcriptional target genes showed the largest normalized number of functional enrichments, compared with target genes including 5 – 60% of randomly selected genes. The normalized number of functional enrichments was changed according to the criteria of enhancer-promoter interactions such as distance from transcriptional start sites and orientation of CTCF-binding sites. Forward-reverse orientation of CTCF-binding sites showed significantly higher normalized number of functional enrichments than the other orientations. Journal papers showed that the top five frequent functional enrichments were related to the cellular functions in the three cell types. The median expression level of transcriptional target genes changed according to the criteria of enhancer-promoter assignments (i.e. interactions) and was correlated with the changes of the normalized number of functional enrichments of transcriptional target genes.

**Conclusions:** Human putative transcriptional target genes showed significant functional enrichments. Functional enrichments were related to the cellular functions. The normalized number of functional enrichments of human putative transcriptional target genes changed according to the criteria of enhancer-promoter assignments and correlated with the median expression level of the target genes. These analyses and characters of human putative transcriptional target genes would be useful to examine the criteria of enhancer-promoter assignments and to predict the novel mechanisms and factors such as DNA binding proteins and DNA sequences of enhancer-promoter interactions.

## Background

More than 400 types of cells have been found in the human body. Human development is accompanied by the differentiation of stem cells into various cell types, leading to a diversification of their phenotypes and functions. For example, the development of the immune system involves differentiation and diversification of stem cells into various types of mature immune cells. The functions of monocytes include phagocytosis and antigen presentation. CD4^+^ T cells, however, play a central role in cell-mediated immunity and are involved in the activation of phagocytes and antigen-specific cytotoxic T-lymphocytes, and the release of various cytokines in response to an antigen. The CD20^+^ B cells are involved in the production of antibodies against antigens.

Differentiation of cells is often triggered by the expression of transcription factors (TF) followed by the expression of their target genes, which results in the transformation of cells into other cell types. For example, the transcription factors PU.1 and CCAAT enhancer-binding protein α (C/EBPα) play a critical role in the expression of myeloid-specific genes and the generation of monocytes and macrophages [1, 2]. The transcription factor GATA-3 is essential for early T cell development and the differentiation of naive CD4^+^ T cells into Th2 effector cells [3]. E2A, EBF1, PAX5, and Ikaros are among the most important transcription factors that control early development in mice, thereby conditioning homeostatic B cell lymphopoiesis [4].

We previously examined the differentiation of monocytes and macrophages in mice, and discovered that the transcription factor IRF8 was essential for cellular differentiation [5]. An analysis of transcription factor-binding sites (TFBS) revealed that IRF8 regulated the expression of KLF4 through the IRF8 transcriptional cascade. Functional enrichment analyses revealed that the target genes of IRF8 showed functional enrichment for antigen presentation, whereas those of KLF4 showed functional enrichments for phagocytosis and locomotion. These results suggested that the transcriptional cascades of IRF8 and KLF4 included different functional modules of target genes.

Functional enrichments of transcriptional cascades of IRF8 and KLF4 appeared to be related to the cellular functions of monocytes and macrophages. Although several transcription factors were expressed in monocytes and macrophages, the number of these transcriptional target genes that resulted in functional enrichments remains unknown. Whether transcriptional target genes in other human cells show functional enrichments remain unclear. If the transcriptional target genes showed significant functional enrichment, analyzing transcriptional target genes would be useful in identifying genes involved in a specific cellular function. Using the budding yeast, previous studies examined the functional enrichments on a genome-scale genetic interaction map using the GeneMANIA algorithm [6-8]. Using bacterial systems, the analyses of functional enrichments of predicted regulatory networks were performed using Gene Ontology annotations [9]. Various databases of functional annotations of genes and pathways exist. Analysis of functional enrichments is expected to be useful for understanding the association of genes involved in similar functions and same pathways, and for predicting unknown gene functions such as non-protein-coding RNAs. In addition, the extent of enhancer contribution to functional enrichments of transcriptional target genes remains unknown.

In this study, transcriptional target genes were predicted using public databases of open chromatin regions of human monocytes, naive CD4^+^ T, CD20^+^ B cells, HUVEC, IMR90, MCF-7, HMEC, H1-hESC, iPSC, and ChIP-seq data of human H1-hESC cells and known transcription factor binding sequences. Functional enrichment analyses of putative transcriptional target genes were conducted using 10 different annotation databases of functional annotations and pathways. The gene expression level of transcriptional target genes was examined in the cells.

## Results

### Prediction of transcriptional target genes

To examine functional enrichments of transcriptional target genes in a genome scale, transcriptional target genes were predicted in human monocytes, CD4^+^ T cells, and CD20^+^ B cells. Searches for known transcription factor binding sequences, which were collected from various databases and papers, were conducted in open chromatin regions of the promoter sequences of RefSeq transcripts (Figure 1, see Methods). Among 6,277 transcription factor binding sequences derived from vertebrates, 4,373 were linked to 1,018 TF transcripts computationally (see Methods). To maintain the sensitivity of the searches for transcription factor binding sites and as some transcription factors will recognize multiple distinctly different sequence motifs, transcription factor binding sequences that targeted the same genes were recognized as redundant, and one of the sequences was used [10] (see Methods). In total, 3,337 transcription factor binding sequences in human monocytes, 3,652 in CD4^+^ T cells, and 3,187 in CD20^+^ B cells were identified with their target genes, which were selected from highly expressed genes in a cell (top 30% expression level, see Methods).

**Figure 1.**
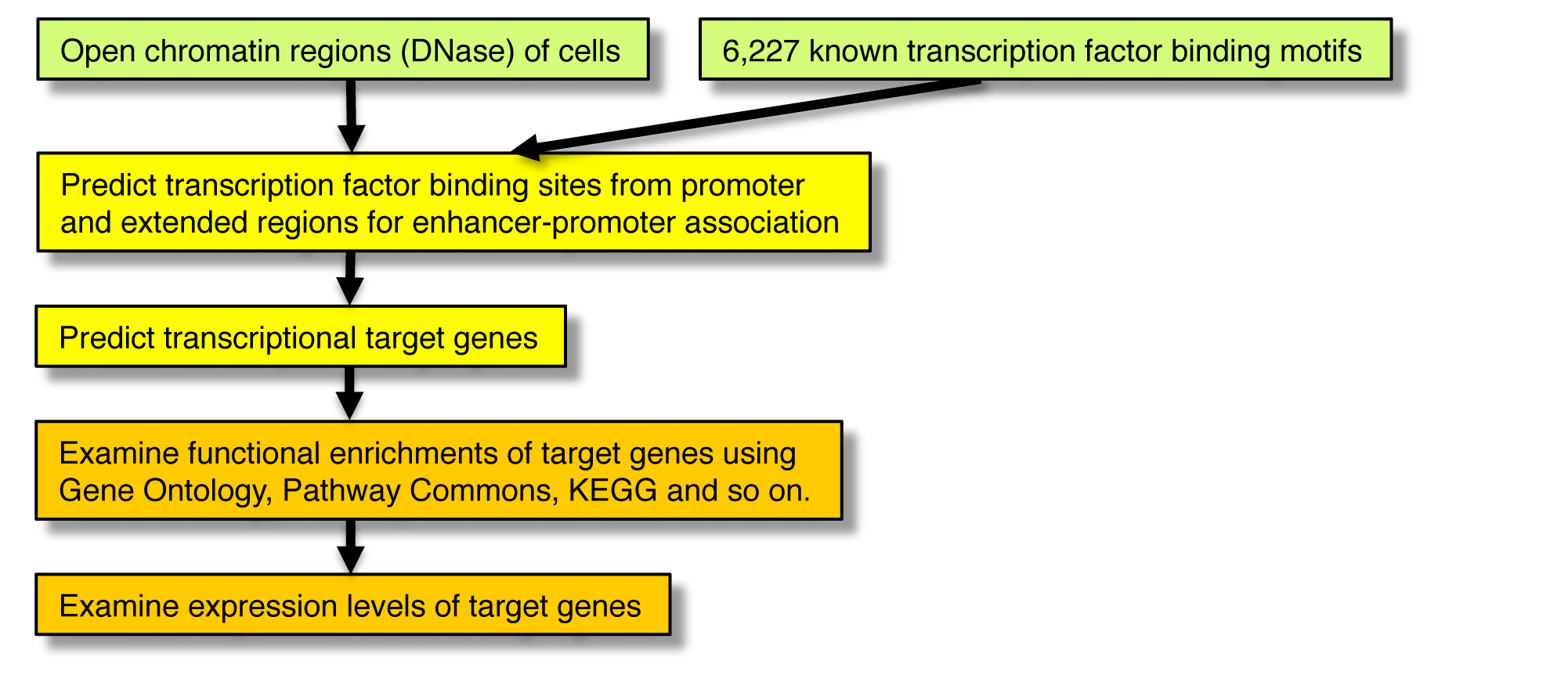
Analyses of functional enrichments of putative transcriptional target genes. Transcriptional target genes were predicted using open chromatin regions (DNase-DGF) and known transcription factor binding sequences. Functional enrichments of target genes were analyzed using 10 annotation databases, and were changed based on the criteria of promoter and extended regions for enhancer-promoter association (EPA). To compare with the tendency of the normalized numbers of functional enrichments, the median expression levels of target genes were examined using promoter and EPA.

The total numbers of unique highly expressed target genes of transcription factor binding sequences were 4,481, 7,558, and 4,753 in monocytes, CD4^+^ T cells, and CD20^+^ B cells respectively using promoters. The mean target genes of a transcription factor were 124, 164, and 144 in monocytes, CD4^+^ T cells, and CD20^+^ B cells, respectively, with the corresponding medians being 24, 33, and 24, respectively. With regard to the genomic localizations of TFBS, 51%, 65%, and 61% of TFBS were located within promoters (±5 kb of TSS) of target genes in monocytes, CD4^+^ T cells, and CD20^+^ B cells, respectively (according to association rule 1, see Methods).

### Functional enrichments of putative transcriptional target genes

Functional enrichments of the putative target genes were examined. The distribution of functional enrichments in transcriptional target genes was predicted using genome sequences of promoters in the three cell types (Figure 1 and Table 1, see Methods). Furthermore, the effect of transcriptional target genes including randomly selected genes on functional enrichments was investigated using DNase-DGF data of monocytes, CD4^+^ T and CD20^+^ B cells, HUVEC, IMR90, MCF-7, HMEC, and ChIP-seq data of H1-hESC (Figure 2A and B, and Figure S1 and S2, see Methods). The native putative transcriptional target genes not including randomly selected genes showed the highest functional enrichments using Gene Ontology, GO Slim, KEGG, Pathway Commons, WikiPathways, InterPro and UniProt functional regions (Domains) in both DNase-DGF and ChIP-seq data of the five types of cells. Of the 10 databases used in this analysis, the Gene Ontology database consists of three types of functional annotations, i.e., 20,836 biological processes, 9,020 molecular functions, and 2,847 cellular components. The numbers of functional enrichments of Gene Ontology annotations in target genes of a transcription factor were 2,902, 4,077, and 2,778 in monocytes, CD4^+^ T cells, and CD20^+^ B cells, respectively. An examination of functional enrichments of highly expressed genes (top 30% expression level), independent of transcriptional target genes, revealed 237, 301, and 239 ‘unique’ Gene Ontology annotations in monocytes, CD4^+^ T cells, and CD20^+^ B cells, respectively (Table 1). Further, the examination of functional enrichments of highly expressed target genes (top 30% expression level) in target genes revealed 1,271, 1,654, and 1,192 ‘unique’ Gene Ontology annotations in monocytes, CD4^+^ T cells, and CD20^+^ B cells, respectively i.e., These numbers were four times larger than functional enrichments identified by gene expression information alone, independent of transcriptional target genes, suggesting that transcriptional target genes were frequently associated with similar functions or pathways (Table S3 and S4).

**Table 1.** Number of functional enrichments and unique functional enrichemnts of putative transcriptional target genes.

| Number of functional enrichments of putative trancriptional target genes |  |  |  |  |  |  |  |  |  |  |
| --- | --- | --- | --- | --- | --- | --- | --- | --- | --- | --- |
|  | KEGG | TF<br>Targets | CTD<br>Ontology | GO<br>Slim | GO | Pathway<br>Commons | BioMarkers | MicroRNA | Domains | WikiPathways |
| Monocyte | 349 | 107 | 209 | 114 | 2,902 | 1,005 | 42 | 451 | 1,202 | 242 |
| CD4<sup>+</sup> T cell | 317 | 135 | 278 | 77 | 4,077 | 1,806 | 47 | 754 | 1,401 | 405 |
| CD20<sup>+</sup> B cell | 323 | 103 | 170 | 88 | 2,778 | 821 | 39 | 948 | 950 | 288 |
| Number of unique functional enrichments of gene expression information alone and putative transcriptional target genes.<br>* Wilcoxon signed-rank test $p < 0.01$ (except for MicroRNA and Domains) | | | | | | | | | | |
| Gene expression information alone |  |  |  |  |  |  |  |  |  |  |
| Monocyte | 43 | 0 | 35 | 11 | 237 | 101 | 7 | 314 | 404 | 58 |
| CD4<sup>+</sup> T cell | 47 | 0 | 19 | 9 | 301 | 165 | 9 | 136 | 397 | 81 |
| CD20<sup>+</sup> B cell | 42 | 0 | 27 | 12 | 239 | 247 | 6 | 370 | 409 | 65 |
| Putative transcriptional target genes |  |  |  |  |  |  |  |  |  |  |
| Monocyte | 95 | 16 | 127 | 12 | 1,271 | 242 | 17 | 97 | 303 | 105 |
| CD4<sup>+</sup> T cell | 105 | 26 | 146 | 23 | 1,654 | 415 | 24 | 224 | 585 | 133 |
| CD20<sup>+</sup> B cell | 93 | 23 | 96 | 23 | 1,192 | 329 | 16 | 231 | 397 | 106 |

**Figure 2.**
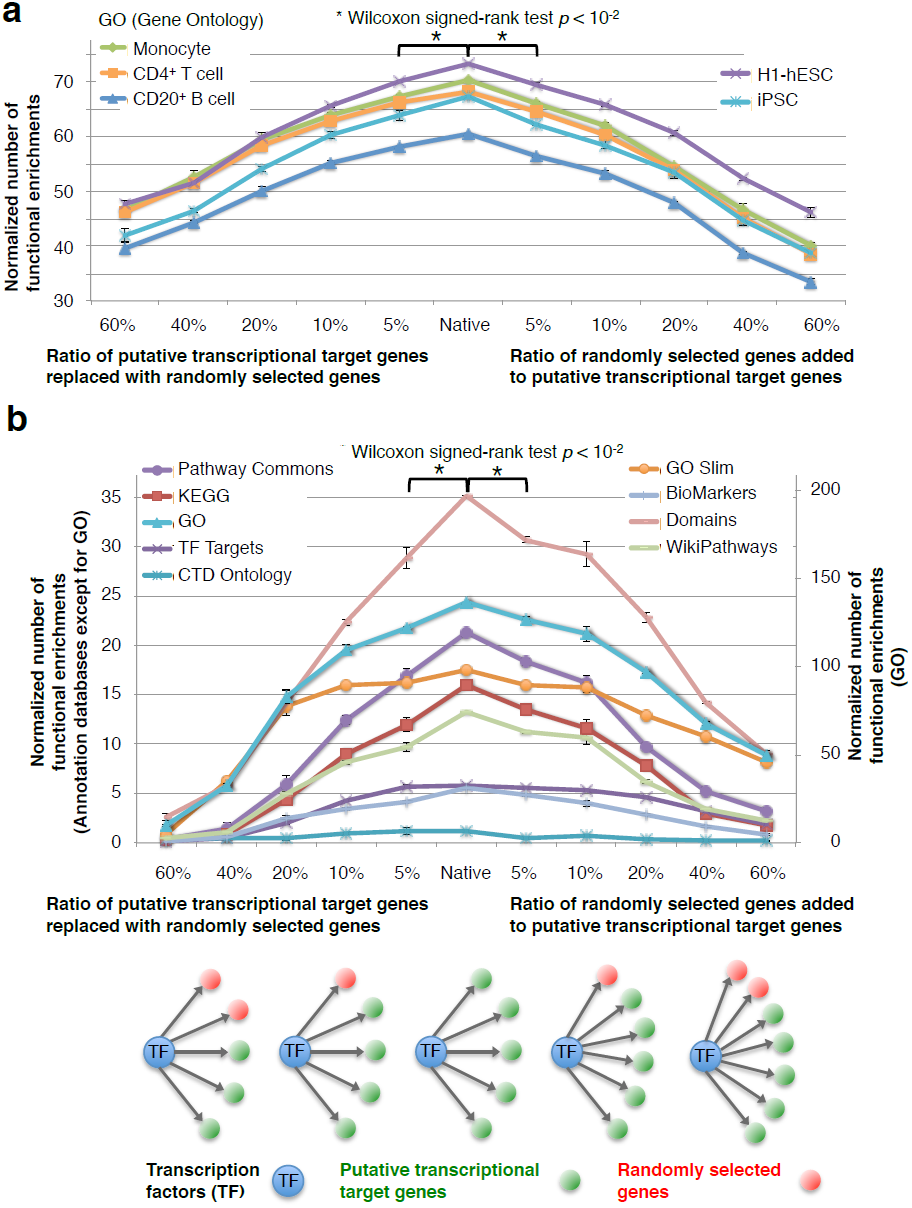
Effect of randomly selected genes on functional enrichments. (**a)** Effect of randomly selected genes on functional enrichments using DNase-DGF data. Transcriptional target genes were predicted using DNase-DGF data in human monocytes, CD4^+^ T, CD20^+^ B, other four somatic and two stem cells (H1-hESC and iPSC) (see also Figure S1). To test whether slight changes of transcriptional target genes were reflected in the normalized number of their functional enrichments, the ratio of randomly selected genes in the target genes of each TF was changed between 5% and 60%. In the left part of the graphs, randomly selected genes were replaced with the target genes where the total number of target genes was unchanged. In the right part of the graphs, randomly selected genes were added to the target genes where the total number of target genes was increased. The result of Gene Ontology annotation was shown. The results of Pathway Commons and KEGG were shown in Figure S1. Native target genes showed the most functional enrichments in most cell types. (**b)** Effect of randomly selected genes on functional enrichments using ChIP-seq data. Transcriptional target genes were predicted using ChIP-seq data of 19 TF in H1-hESC. The results of nine functional annotation databases were shown and the result of target genes of microRNAs was shown in Figure S2. Native target genes showed the most functional enrichments using most annotation databases except for low frequent functional annotations. Putative transcriptional target genes tend to include similar function of genes.

Functional enrichments of transcriptional target genes from other databases were also examined (Table 1). KEGG, Target genes of transcription factors, Disease Ontology, GO Slim, Pathway Commons, Cellular biomarkers, Target genes of microRNAs, Protein domains, and WikiPathways had 95, 16, 127, 12, 242, 17, 97, 303, and 105 unique functional annotations, respectively. The numbers of functional enrichments of transcriptional target genes in the other annotation databases except for microRNAs and Protein domains were significantly higher than gene expression information alone, independent of transcriptional target genes, as well as Gene Ontology annotations (Table 1). The functional enrichments of transcriptional target genes from Pathway Commons for monocytes, CD4^+^ T cells, and CD20^+^ B cells are shown in Table 2 and Table S5. Functional enrichments were found to be related to cellular functions, e.g., interferon signaling, GMCSF (Granulocyte-macrophage colony-stimulating factor, a kind of cytokine)-mediated signaling events, antigen processing-cross presentation in monocytes; TCR (T-cell receptor) signaling in naive CD4^+^ T cells, IL-12 (Interleukin-12, a kind of cytokine)-mediated signaling events, and downstream signaling in naive CD8^+^ T cells in CD4^+^ T cells; interferon alpha/beta signaling, IL8- and CXCR2 (Chemokine receptor type 2, a kind of cytokine)-mediated signaling events, and BCR (B cell antigen receptor) signaling pathway in CD20^+^ B cells. WikiPathways, KEGG and GO also revealed that functional enrichments were associated with cellular functions (Table S6, S7 and S8).

**Table 2.** Functional enrichments of putative transcriptional target genes using Pathway Commons.

| Monocyte - Pathway Commons | No. of TFs |
| --- | --- |
| Proteoglycan syndecan-mediated signaling events | 18 |
| Regulation of CDC42 activity | 15 |
| LKB1 signaling events | 14 |
| Glypican pathway | 13 |
| Interferon Signaling | 13 |
| Sphingosine 1-phosphate (S1P) pathway | 13 |
| IL5-mediated signaling events | 12 |
| Syndecan-1-mediated signaling events | 12 |
| IL3-mediated signaling events | 11 |
| Mitotic Prophase | 11 |
| Golgi Cisternae Pericentriolar Stack Reorganization | 11 |
| Interferon alpha/beta signaling | 11 |
| IFN-gamma pathway | 10 |
| Signaling events mediated by Hepatocyte Growth Factor Receptor (c-Met) | 10 |
| Recruitment of mitotic centrosome proteins and complexes | 10 |
| CD4<sup>+</sup> T cell - Pathway Commons | No. of TFs |
| TCR signaling in naive CD8<sup>+</sup> T cells | 36 |
| IL12-mediated signaling events | 24 |
| Downstream signaling in naive CD8<sup>+</sup> T cells | 21 |
| TCR signaling in naive CD4<sup>+</sup> T cells | 21 |
| IL12 signaling mediated by STAT4 | 20 |
| Validated transcriptional targets of AP1 family members Fra1 and Fra2 | 19 |
| CXCR4-mediated signaling events | 17 |
| ATF-2 transcription factor network | 15 |
| Thrombin/protease-activated receptor (PAR) pathway | 14 |
| TCR signaling | 14 |
| PAR1-mediated thrombin signaling events | 14 |
| Downstream TCR signaling | 14 |
| Internalization of ErbB1 | 13 |
| Urokinase-type plasminogen activator (uPA) and uPAR-mediated signaling | 13 |
| ErbB receptor signaling network | 13 |
| CD20<sup>+</sup> B cell - Pathway Commons | No. of TFs |
| Interferon alpha/beta signaling | 12 |
| Alpha6Beta4Integrin | 11 |
| Validated targets of C-MYC transcriptional activation | 11 |
| IL8- and CXCR2-mediated signaling events | 10 |
| Antigen processing-Cross presentation | 9 |
| BCR signaling pathway | 9 |
| IL6-mediated signaling events | 9 |
| Cell junction organization | 9 |
| ER-Phagosome pathway | 8 |
| Regulation of CDC42 activity | 8 |
| CXCR4-mediated signaling events | 8 |
| CDC42 signaling events | 8 |
| Syndecan-4-mediated signaling events | 8 |
| Noncanonical Wnt signaling pathway | 7 |
| Class I MHC mediated antigen processing & presentation | 7 |

### Effect of enhancer-promoter association rules on functional enrichments

To understand the effect of ‘promoter and extended regions for enhancer-promoter association (EPA)’ on the functional enrichments of target genes, the rule of extended regions was modified according to four criteria (Figure 3A and see Methods) [11], and functional enrichments were investigated.

**Figure 3.**
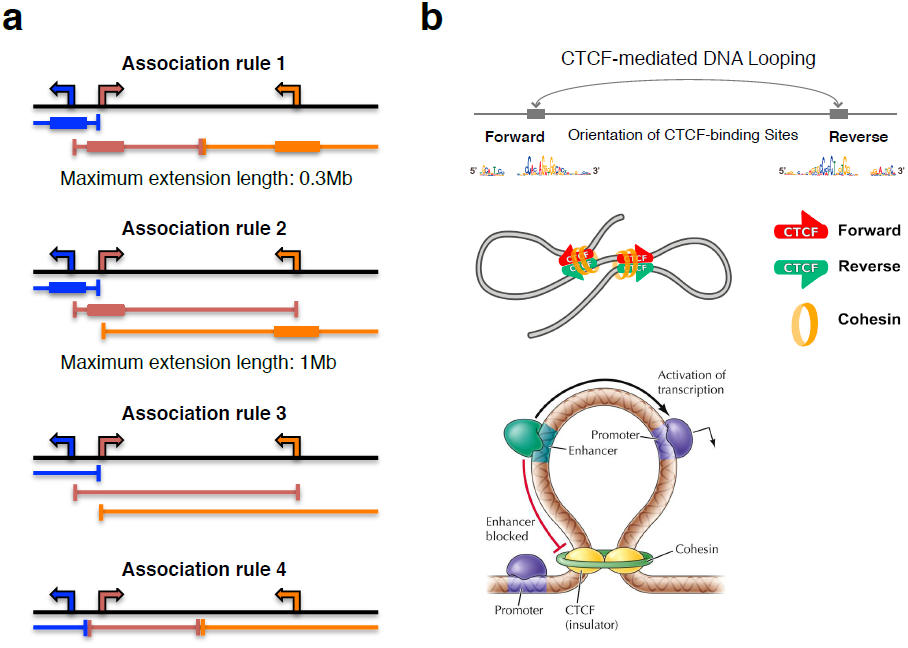
Criteria of promoter and extended regions for enhancer-promoter association and features of chromatin interactions. (**a**) Computationally-defined regulatory domains [11]. The transcription start site (TSS) of each gene is indicated as an arrow. The corresponding regulatory domain for each gene is shown in a matching color as a bracketed line. The *basal plus extension* association rule assigns a basal regulatory domain to each gene regardless of genes nearby (thick line, Association rule 1 and 2) (see Methods). The domain is then extended to the basal regulatory domain of the nearest upstream and downstream genes. The *two nearest genes* association rule extends the regulatory domain to the TSS of the nearest upstream and downstream genes (Association rule 3). The *single nearest gene* association rule extends the.regulatory domain to the midpoint between this gene’s TSS and the nearest gene’s TSS both upstream and downstream (Association rule 4). (**b**) Forward–reverse orientation of CTCF-binding sites are frequently found in chromatin interactions. CTCF can block the interaction between enhancers and promoters limiting the activity of enhancers to certain functional domains. Figures adapted from [13, 69] and the bottom figure in panel b was in the book ‘THE CELL: A molecular approach 6e, Figure 7.21, 2013 Sinauer Associates, Inc’ and Image 68 insulators in ‘http://slideplayer.com/slide/3836520’.

According to the association rule (1), the means of target genes were 177, 217, and 175 in monocytes, CD4^+^ T cells, and CD20^+^ B cells, respectively, whereas the corresponding medians were 55, 58, and 37, respectively (Table S9). The numbers of functional enrichments of Pathway Commons annotations using promoter regions were 1,005, 1,806, and 821 in monocytes, CD4^+^ T cells, and CD20^+^ B cells, respectively (Table S10). With the use of EPA (association rule 1), the numbers of functional enrichments of Pathway Commons annotations were 3,087, 7,216, and 3,900, representing 3.07-, 4.00-, and 4.75-fold increases, respectively, in the three cells types. Additionally, the numbers of ‘unique’ Pathway Commons annotations with promoter regions were 321, 415, and 329 in monocytes, CD4^+^ T cells, and CD20^+^ B cells, respectively; the corresponding numbers with the use of EPA (association rule 1) were 364, 437, and 364, representing 1.13-, 1.05-, and 1.11-fold increases, respectively, in the three cell types. The normalized numbers of functional enrichments of Pathway Commons annotations were 44.75, 84.51, and 59.32, representing 1.84-, 2.80-, and 3.32-fold increases, respectively, in the three cell types (association rule 1, Table 3). Other cell types also showed the same tendencies (Table 3).

**Table 3.**
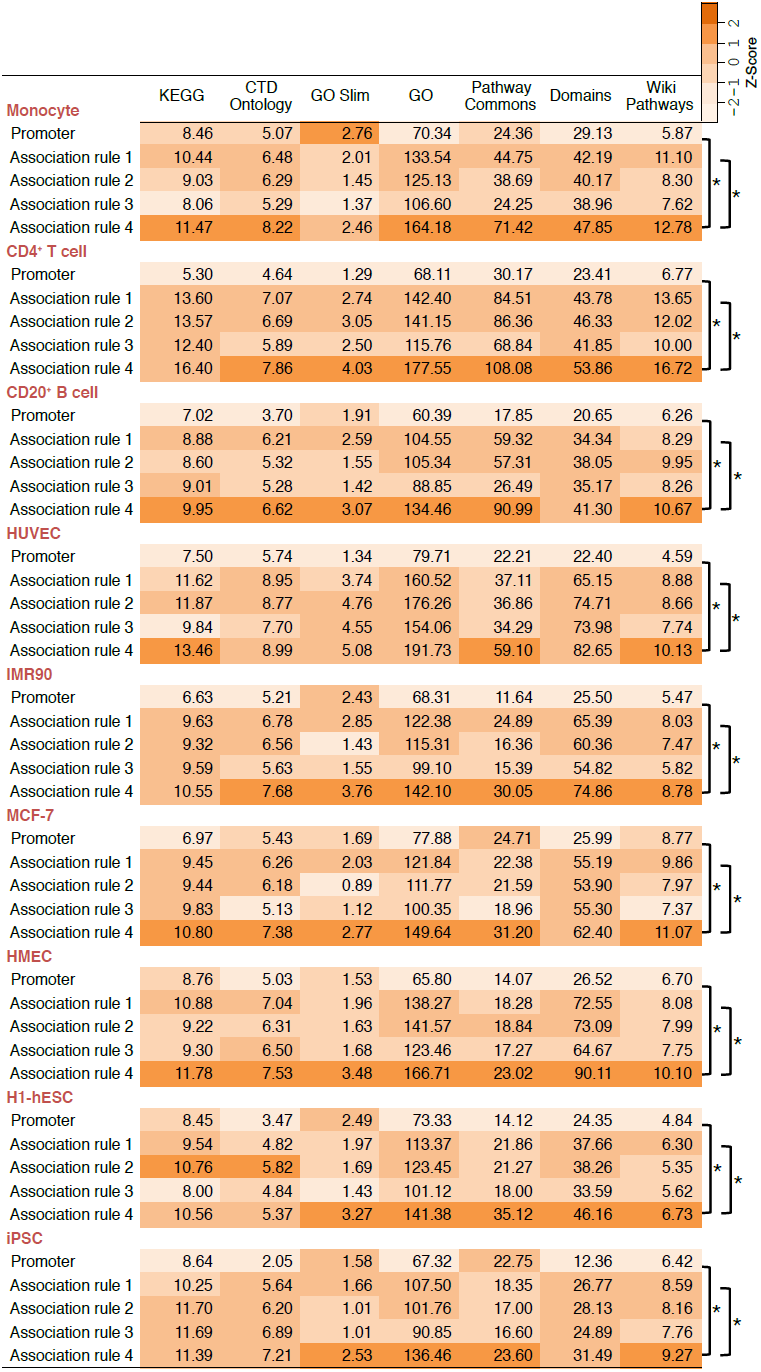
Normalized number of functional enrichments of putative transcriptional target genes using promoter and extended regions for enhancer-promoter association. * Wilcoxon signed-rank test *p* < 0.05.

|  | KEGG | CTD<br>Ontology | GO Slim | GO | Pathway<br>Commons | Domains | Wiki<br>Pathways | Z-Score |
| --- | --- | --- | --- | --- | --- | --- | --- | --- |
| **Monocyte** |  |  |  |  |  |  |  |  |
| Promoter | 8.46 | 5.07 | 2.76 | 70.34 | 24.36 | 29.13 | 5.87 |  |
| Association rule 1 | 10.44 | 6.48 | 2.01 | 133.54 | 44.75 | 42.19 | 11.10 | *<br>* |
| Association rule 2 | 9.03 | 6.29 | 1.45 | 125.13 | 38.69 | 40.17 | 8.30 |  |
| Association rule 3 | 8.06 | 5.29 | 1.37 | 106.60 | 24.25 | 38.96 | 7.62 |  |
| Association rule 4 | 11.47 | 8.22 | 2.46 | 164.18 | 71.42 | 47.85 | 12.78 |  |
| **CD4<sup>+</sup> T cell** |  |  |  |  |  |  |  |  |
| Promoter | 5.30 | 4.64 | 1.29 | 68.11 | 30.17 | 23.41 | 6.77 |  |
| Association rule 1 | 13.60 | 7.07 | 2.74 | 142.40 | 84.51 | 43.78 | 13.65 | *<br>* |
| Association rule 2 | 13.57 | 6.69 | 3.05 | 141.15 | 86.36 | 46.33 | 12.02 |  |
| Association rule 3 | 12.40 | 5.89 | 2.50 | 115.76 | 68.84 | 41.85 | 10.00 |  |
| Association rule 4 | 16.40 | 7.86 | 4.03 | 177.55 | 108.08 | 53.86 | 16.72 |  |
| **CD20<sup>+</sup> B cell** |  |  |  |  |  |  |  |  |
| Promoter | 7.02 | 3.70 | 1.91 | 60.39 | 17.85 | 20.65 | 6.26 |  |
| Association rule 1 | 8.88 | 6.21 | 2.59 | 104.55 | 59.32 | 34.34 | 8.29 | *<br>* |
| Association rule 2 | 8.60 | 5.32 | 1.55 | 105.34 | 57.31 | 38.05 | 9.95 |  |
| Association rule 3 | 9.01 | 5.28 | 1.42 | 88.85 | 26.49 | 35.17 | 8.26 |  |
| Association rule 4 | 9.95 | 6.62 | 3.07 | 134.46 | 90.99 | 41.30 | 10.67 |  |
| **HUVEC** |  |  |  |  |  |  |  |  |
| Promoter | 7.50 | 5.74 | 1.34 | 79.71 | 22.21 | 22.40 | 4.59 |  |
| Association rule 1 | 11.62 | 8.95 | 3.74 | 160.52 | 37.11 | 65.15 | 8.88 | *<br>* |
| Association rule 2 | 11.87 | 8.77 | 4.76 | 176.26 | 36.86 | 74.71 | 8.66 |  |
| Association rule 3 | 9.84 | 7.70 | 4.55 | 154.06 | 34.29 | 73.98 | 7.74 |  |
| Association rule 4 | 13.46 | 8.99 | 5.08 | 191.73 | 59.10 | 82.65 | 10.13 |  |
| **IMR90** |  |  |  |  |  |  |  |  |
| Promoter | 6.63 | 5.21 | 2.43 | 68.31 | 11.64 | 25.50 | 5.47 |  |
| Association rule 1 | 9.63 | 6.78 | 2.85 | 122.38 | 24.89 | 65.39 | 8.03 | *<br>* |
| Association rule 2 | 9.32 | 6.56 | 1.43 | 115.31 | 16.36 | 60.36 | 7.47 |  |
| Association rule 3 | 9.59 | 5.63 | 1.55 | 99.10 | 15.39 | 54.82 | 5.82 |  |
| Association rule 4 | 10.55 | 7.68 | 3.76 | 142.10 | 30.05 | 74.86 | 8.78 |  |
| **MCF-7** |  |  |  |  |  |  |  |  |
| Promoter | 6.97 | 5.43 | 1.69 | 77.88 | 24.71 | 25.99 | 8.77 |  |
| Association rule 1 | 9.45 | 6.26 | 2.03 | 121.84 | 22.38 | 55.19 | 9.86 | *<br>* |
| Association rule 2 | 9.44 | 6.18 | 0.89 | 111.77 | 21.59 | 53.90 | 7.97 |  |
| Association rule 3 | 9.83 | 5.13 | 1.12 | 100.35 | 18.96 | 55.30 | 7.37 |  |
| Association rule 4 | 10.80 | 7.38 | 2.77 | 149.64 | 31.20 | 62.40 | 11.07 |  |
| **HMEC** |  |  |  |  |  |  |  |  |
| Promoter | 8.76 | 5.03 | 1.53 | 65.80 | 14.07 | 26.52 | 6.70 |  |
| Association rule 1 | 10.88 | 7.04 | 1.96 | 138.27 | 18.28 | 72.55 | 8.08 | *<br>* |
| Association rule 2 | 9.22 | 6.31 | 1.63 | 141.57 | 18.84 | 73.09 | 7.99 |  |
| Association rule 3 | 9.30 | 6.50 | 1.68 | 123.46 | 17.27 | 64.67 | 7.75 |  |
| Association rule 4 | 11.78 | 7.53 | 3.48 | 166.71 | 23.02 | 90.11 | 10.10 |  |
| **H1-hESC** |  |  |  |  |  |  |  |  |
| Promoter | 8.45 | 3.47 | 2.49 | 73.33 | 14.12 | 24.35 | 4.84 |  |
| Association rule 1 | 9.54 | 4.82 | 1.97 | 113.37 | 21.86 | 37.66 | 6.30 | *<br>* |
| Association rule 2 | 10.76 | 5.82 | 1.69 | 123.45 | 21.27 | 38.26 | 5.35 |  |
| Association rule 3 | 8.00 | 4.84 | 1.43 | 101.12 | 18.00 | 33.59 | 5.62 |  |
| Association rule 4 | 10.56 | 5.37 | 3.27 | 141.38 | 35.12 | 46.16 | 6.73 |  |
| **iPSC** |  |  |  |  |  |  |  |  |
| Promoter | 8.64 | 2.05 | 1.58 | 67.32 | 22.75 | 12.36 | 6.42 |  |
| Association rule 1 | 10.25 | 5.64 | 1.66 | 107.50 | 18.35 | 26.77 | 8.59 | *<br>* |
| Association rule 2 | 11.70 | 6.20 | 1.01 | 101.76 | 17.00 | 28.13 | 8.16 |  |
| Association rule 3 | 11.69 | 6.89 | 1.01 | 90.85 | 16.60 | 24.89 | 7.76 |  |
| Association rule 4 | 11.39 | 7.21 | 2.53 | 136.46 | 23.60 | 31.49 | 9.27 |  |

The normalized numbers of the functional enrichments of transcriptional target genes showed association rule (4) as the highest number, followed by association rule (1) and (2) in the three cell types. Although association rule (3) was the longest among the four criteria, it showed the lowest number of functional enrichments in the three cell types (Figure 3A and Table 3). ChIP-seq data of 19 TF in H1-hESC (Human embryonic stem cells) also showed almost the same tendency (difference between association rule (4) and (1) was not statistically significant, probably due to a large number of transcriptional target genes predicted using 19 TF ChIP-seq data. Several thousands of target genes of each TF were predicted. Some of them would be indirect interactions between TF and genome DNA, which were identified by ChIP-seq experiments. (Table S11, see Additional file).

Differences in functional enrichments using Pathway Commons were examined between promoters *versus* EPA (association rule 1) (Table 4 and Table S12). A comparison of 321 and 364 functional enrichments using the promoters and EPA, respectively, in monocytes revealed that 152 (47% in promoters, 42% in extended regions) of them were common. For example, IFN-gamma (Interferon gamma) pathway, GMCSF (Granulocyte-macrophage colony-stimulating factor, a kind of cytokine)-mediated signaling events, and PDGF (Platelet-derived growth factor) receptor signaling network were enriched using extended regions (association rule 1) as opposed to promoters (Table S12). The comparison of 415 (promoters) and 437 (extended regions) functional enrichments in CD4^+^ T cells revealed that 163 of them (39% in promoters, 37% in extended regions) were common. IFN-gamma pathway, TCR (T-cell receptor) signaling in naive CD4^+^ T cells, and IL3 (Interleukin-3, a kind of cytokine)-mediated signaling events were enriched using extended regions. The comparison of 329 (promoters) and 364 (extended regions) functional enrichments in CD20^+^ B cells revealed that 171 of them (52% in promoters, 47% in extended regions) were common. IL5-mediated signaling events, IL4-mediated signaling events, and cytokine signaling in immune system were enriched in CD20^+^ B cells using extended regions. Only about 40% of functional enrichments of Pathway Commons annotations were unchanged between promoters and EPA. EPA significantly affected the functional enrichments of transcriptional target genes. Journal papers showed that frequent functional enrichments were related to the cellular functions in the three cell types (Table 4). These results showed that new functional enrichments related to cellular functions were identified using extended regions for enhancer-promoter association.

**Table 4.** Differences in functional enrichments using Pathway Commons. Immune cell-related functional annotations were enriched more in ‘promoter and extended regions for enhancer-promoter association (EPA)’ than promoters. Five annotations are shown in each cell type. These functions are confirmed to be related to the cellular functions by reference journal papers.

| Cell type | Term of functional annotation | No. of TF<br>EPA (Association rule 4) | No. of TF<br>Promoter | P-value | Experimental<br>support |
| --- | --- | --- | --- | --- | --- |
| Monocytes | IFN-gamma pathway | 51 | 10 | $6.89 \times 10^{-4}$ | Ref. 39 |
| | IL3-mediated signaling events | 46 | 11 | $4.69 \times 10^{-3}$ | Ref. 40 |
| | GMCSF-mediated signaling events | 43 | 10 | $5.25 \times 10^{-3}$ | Ref. 41 |
| | PDGF receptor signaling network | 43 | 8 | $1.34 \times 10^{-2}$ | Ref. 42 |
| | VEGF and VEGFR signaling network | 44 | 9 | $2.10 \times 10^{-3}$ | Ref. 43 |
| CD4<sup>+</sup> T cell | Integrin family cell surface interactions | 112 | 13 | $2.59 \times 10^{-12}$ | Ref. 44 |
| | IFN-gamma pathway | 107 | 12 | $5.26 \times 10^{-12}$ | Ref. 45 |
| | LKB1 signaling events | 107 | 11 | $1.96 \times 10^{-12}$ | Ref. 46 |
| | TCR signaling in naive CD4<sup>+</sup> T cells | 103 | 21 | $3.98 \times 10^{-8}$ | Ref. 47 |
| | IL3-mediated signaling events | 100 | 8 | $9.54 \times 10^{-13}$ | Ref. 48 |
| CD20<sup>+</sup> B cell | Integrin family cell surface interactions | 69 | 2 | $5.17 \times 10^{-11}$ | Ref. 49 |
| | IL5-mediated signaling events | 64 | 0 | $2.21 \times 10^{-11}$ | Ref. 50 |
| | Insulin Pathway | 63 | 0 | $3.15 \times 10^{-11}$ | Ref. 51 |
| | mTOR signaling pathway | 63 | 0 | $3.15 \times 10^{-11}$ | Ref. 52 |
| | Sphingosine 1-phosphate (S1P) pathway | 63 | 0 | $3.15 \times 10^{-11}$ | Ref. 53 |

Chi-square tests were conducted using the total number of putative transcriptional target genes (Table S9).

### Effect of CTCF-binding sites on functional enrichments

CTCF have the activity of insulators to block the interaction between enhancers and promoters [12]. Recent studies identified a correlation between the orientation of CTCF-binding sites and chromatin loops (Figure 3B) [13]. Forward–reverse (FR) orientation of CTCF-binding sites are frequently found in chromatin loops. To examine the effect of forward–reverse orientation of CTCF-binding sites on functional enrichments of target genes, ‘promoter and extended regions for enhancer-promoter association (EPA)’ were shortened at the genomic locations of forward–reverse orientation of CTCF-binding sites, and transcriptional target genes were predicted from the shortened regions using TFBS (see Methods). The numbers of functional enrichments of target genes were investigated. According to EPA (association rule 4) that were shortened at genomic locations of forward–reverse orientation of CTCF-binding sites, the means of target genes were 67, 64, and 77 in monocytes, CD4^+^ T cells, and CD20^+^ B cells, respectively, whereas the corresponding medians were 23, 21, and 20, respectively (Table S13). The normalized numbers of functional enrichments of Pathway Commons annotations using EPA were 71.42, 108.08, and 90.99 in monocytes, CD4^+^ T cells, and CD20^+^ B cells, respectively (Table 5). With the use of EPA shortened at forward–reverse orientation of CTCF-binding sites, the normalized numbers of functional enrichments of Pathway Commons annotations were 196.58, 220.54, and 220.77, representing 2.75-, 2.04-, and 2.43-fold increases, respectively, in the three cells types. Additionally, the normalized numbers of functional enrichments of ‘unique’ Pathway Commons annotations with EPA were 5.09, 5.34, and 6.00 in monocytes, CD4^+^ T cells, and CD20^+^ B cells, respectively; the corresponding normalized numbers with the use of EPA shortened at forward–reverse orientation of CTCF-binding sites were 9.88, 10.72, and 9.10, representing 1.94-, 2.01-, and 1.52-fold increases, respectively, in the three cell types (Table S14). Other cell types also showed the same tendencies (Table 5). The normalized numbers of functional enrichments were significantly increased between EPA and EPA shortened at forward–reverse orientation of CTCF-binding sites in Gene Ontology, Disease Ontology, Pathway Commons, GO Slim, WikiPathways, KEGG, InterPro and UniProt functional regions (Domains) annotations. These increases were also significant, compared with EPA shortened at CTCF-binding sites without the consideration of their orientation. Transcriptional target genes predicted from EPA shortened at forward–reverse orientation of CTCF-binding sites tend to include similar function of genes significantly.

**Table 5.** Normalized number of functional enrichments of putative transcriptional target genes using CTCF binding sites. ^*^ Wilcoxon signed-rank test *p* < 0.05.

|  | KEGG | CTD<br>Ontology | GO Slim | GO | Pathway<br>Commons | Domains | Wiki<br>Pathways | Z-Score |
| --- | --- | --- | --- | --- | --- | --- | --- | --- |
| **Monocytes** |  |  |  |  |  |  |  |  |
| Association rule 4 | 11.47 | 8.22 | 2.46 | 164.18 | 71.42 | 47.85 | 12.78 | *<br>* |
| CTCF (FR+RF+FF+RR) | 13.19 | 10.39 | 2.74 | 134.26 | 34.37 | 44.46 | 12.96 |  |
| CTCF (FR) | 42.92 | 19.53 | 5.66 | 509.86 | 196.58 | 112.14 | 35.11 |  |
| **CD4<sup>+</sup> T cells** |  |  |  |  |  |  |  |  |
| Association rule 4 | 16.40 | 7.86 | 4.03 | 177.55 | 108.08 | 53.86 | 16.72 | *<br>*<br>* |
| CTCF (FR+RF+FF+RR) | 26.33 | 8.73 | 5.11 | 206.05 | 130.71 | 57.53 | 23.56 |  |
| CTCF (FR) | 69.39 | 14.66 | 24.91 | 560.44 | 220.54 | 133.26 | 46.54 |  |
| **CD20<sup>+</sup> B cells** |  |  |  |  |  |  |  |  |
| Association rule 4 | 9.95 | 6.62 | 3.07 | 134.46 | 90.99 | 41.30 | 10.67 | *<br>*<br>* |
| CTCF (FR+RF+FF+RR) | 8.78 | 4.22 | 2.78 | 94.13 | 27.70 | 28.55 | 7.08 |  |
| CTCF (FR) | 28.86 | 9.72 | 6.61 | 304.01 | 220.77 | 99.89 | 22.68 |  |
| **HUVEC** |  |  |  |  |  |  |  |  |
| Association rule 4 | 13.46 | 8.99 | 5.08 | 191.73 | 59.10 | 82.65 | 10.13 | *<br>*<br>* |
| CTCF (FR+RF+FF+RR) | 19.35 | 10.59 | 3.85 | 155.11 | 28.36 | 57.87 | 13.58 |  |
| CTCF (FR) | 33.71 | 14.82 | 16.33 | 454.47 | 61.85 | 163.38 | 29.38 |  |
| **IMR90** |  |  |  |  |  |  |  |  |
| Association rule 4 | 10.55 | 7.68 | 3.76 | 142.10 | 30.05 | 74.86 | 8.78 | *<br>*<br>* |
| CTCF (FR+RF+FF+RR) | 13.88 | 17.55 | 9.22 | 252.24 | 39.92 | 123.18 | 22.45 |  |
| CTCF (FR) | 30.22 | 15.89 | 11.34 | 402.40 | 53.56 | 180.71 | 37.42 |  |
| **MCF-7** |  |  |  |  |  |  |  |  |
| Association rule 4 | 10.80 | 7.38 | 2.77 | 149.64 | 31.20 | 62.40 | 11.07 | *<br>*<br>* |
| CTCF (FR+RF+FF+RR) | 11.39 | 8.30 | 4.92 | 234.48 | 30.40 | 126.70 | 13.99 |  |
| CTCF (FR) | 22.34 | 11.48 | 6.09 | 397.69 | 51.39 | 145.43 | 37.58 |  |
| **HMEC** |  |  |  |  |  |  |  |  |
| Association rule 4 | 11.78 | 7.53 | 3.48 | 166.71 | 23.02 | 90.11 | 10.10 | *<br>*<br>* |
| CTCF (FR+RF+FF+RR) | 19.68 | 13.54 | 4.29 | 232.47 | 27.97 | 106.24 | 9.84 |  |
| CTCF (FR) | 29.59 | 14.46 | 6.97 | 390.45 | 43.90 | 124.81 | 30.78 |  |
| **H1-hESC** |  |  |  |  |  |  |  |  |
| Association rule 4 | 10.56 | 5.37 | 3.27 | 141.38 | 35.12 | 46.16 | 6.73 | *<br>*<br>* |
| CTCF (FR+RF+FF+RR) | 12.95 | 5.37 | 2.45 | 131.46 | 29.57 | 44.15 | 11.08 |  |
| CTCF (FR) | 28.21 | 10.55 | 9.02 | 303.41 | 56.95 | 139.44 | 19.72 |  |
| **iPSC** |  |  |  |  |  |  |  |  |
| Association rule 4 | 11.39 | 7.21 | 2.53 | 136.46 | 23.60 | 31.49 | 9.27 | *<br>*<br>* |
| CTCF (FR+RF+FF+RR) | 17.17 | 2.33 | 1.69 | 120.87 | 25.60 | 21.06 | 6.42 |  |
| CTCF (FR) | 36.94 | 6.09 | 6.86 | 274.30 | 31.37 | 83.40 | 16.80 |  |

Differences in functional enrichments obtained using EPA *versus* EPA shortened at forward–reverse orientation of CTCF-binding sites were examined using the functional enrichments of Pathway Commons (Table 6 and Results in Additional file). Transcriptional target genes predicted from EPA shortened at the CTCF-binding sites tended to include the similar function of genes. About 40 – 80% of functional enrichments were unchanged between promoters and EPA shortened at forward–reverse orientation of CTCF-binding sites, and the functional enrichments observed in EPA shortened at forward–reverse orientation of CTCF-binding sites as opposed to promoters included various immunological terms. Journal papers showed that the top five frequent functional enrichments were related to the cellular functions in the three cell types (Table 6). These results showed that new functional enrichments related to cellular functions were identified using forward–reverse orientation of CTCF-binding sites.

**Table 6.**
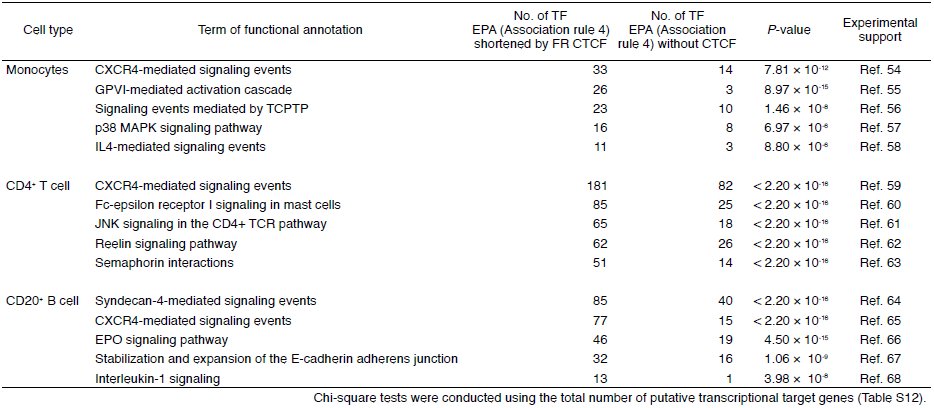
Differences in functional enrichments using Pathway Commons. Immune cell-related functional annotations were enriched more in EPA shortened at forward–reverse orientation of CTCF-binding sites than EPA without CTCF. Five annotations are shown in each cell type. These functions are confirmed to be related to the cellular functions by reference journal papers.

### Comparison of expression levels of putative transcriptional target genes

To examine the relationship between functional enrichments and expression levels of target genes, the expression levels of target genes predicted from promoters and three types of ‘promoter and extended regions for enhancer-promoter assignment (EPA)’ were investigated in monocytes, CD4^+^ T, H1-hESC and iPSC (Figure 4). Median expression levels of the target genes of the same transcription factor binding sequences were compared between promoters and three types of EPA. Red and blue dots in Figure 4 show statistically significant difference of the distribution of expression levels of target genes between promoters and EPA. Additionally, “red dots” show the median expression level of target genes of a TFBS was ‘higher’ in EPA than promoters, and “blue dots” show the median expression level of target genes of a TFBS was ‘lower’ in EPA than promoters. The ratios of red dots were higher in EPA (association rule 4) that were shortened at forward–reverse orientation of CTCF-binding sites *versus* promoters (left graph in Figure 4) than EPA (association rule 4) *versus* promoters (right graph) in monocytes and CD4^+^ T cells. The ratios of blue dots were higher in EPA (association rule 4) that were shortened at forward–reverse orientation of CTCF-binding sites *versus* promoters (left graph) than EPA (association rule 4) *versus* promoters (right graph) in H1-hESC and iPSC. Moreover, the ratio of the sum of median expression levels between the three types of EPA and promoters in monocytes and CD4^+^ T cells was the highest in EPA shortened at forward–reverse orientation of CTCF-binding sites (Table S16). Conversely, the ratio of the sum of median expression levels between the three types of EPA and promoters in H1-hESC and iPSC was the lowest in EPA shortened at forward–reverse orientation of CTCF-binding sites.

**Figure 4.**
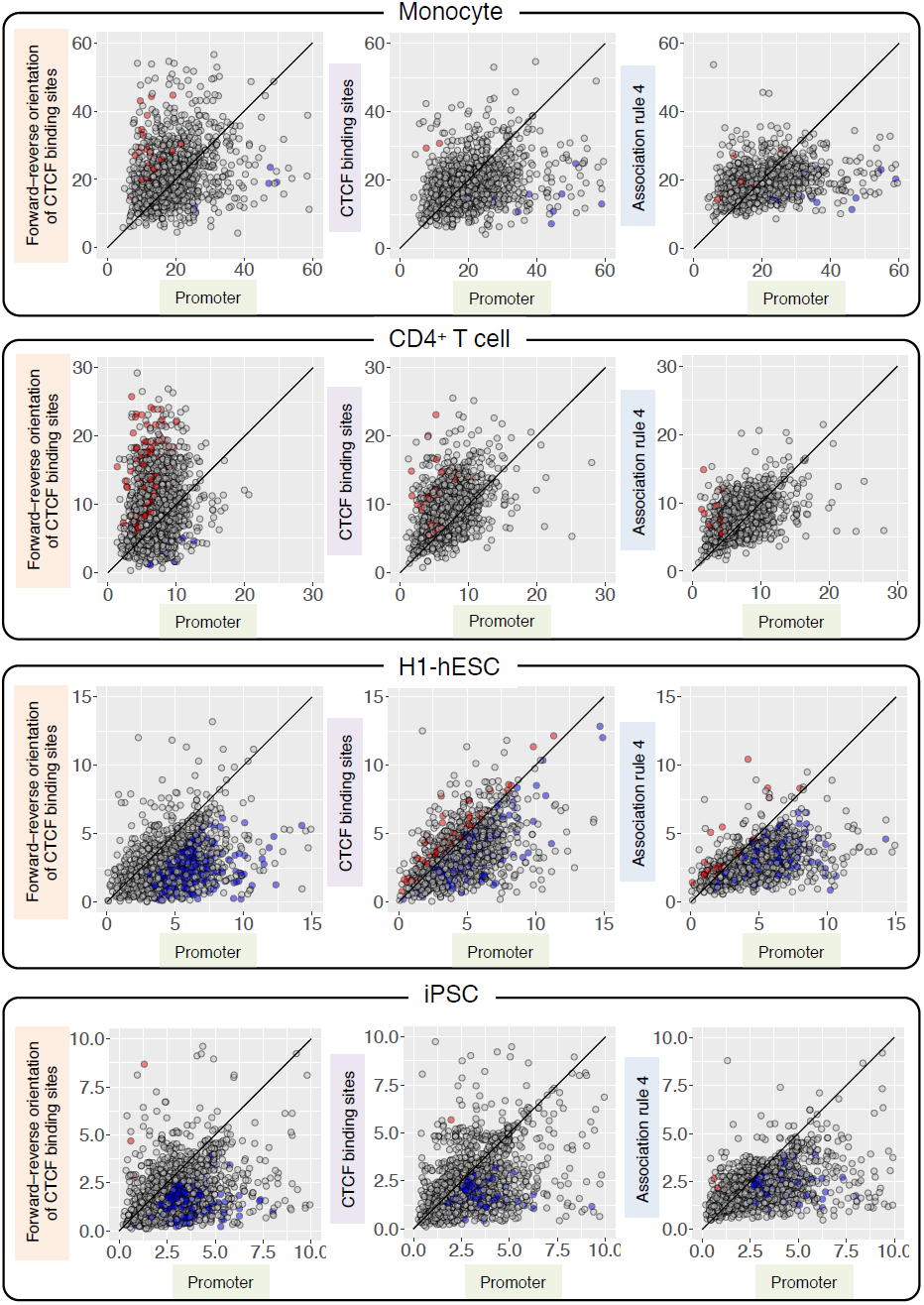
Comparison of the median expression levels of transcriptional target genes. The median expression levels of the target genes of the same transcription factor binding sequences were compared between promoters and three types of promoter and extended regions for enhancer-promoter association (EPA). Red and blue dots show statistically significant difference of the distribution of expression levels of target genes between promoters and EPA. Red dots show the median expression level of target genes was higher in EPA than promoters, and blue dots show the median expression level of target genes was lower in EPA than promoters. The median expression levels of putative transcriptional target genes were significantly lower using TF bound in enhancers than using TF bound in promoters in stem cells such as H1-hESC and iPSC, and higher by TF bound in enhancers in immune cells. These results implied that enhancers significantly affect the expression of target genes.

EPA shortened at forward–reverse orientation of CTCF-binding sites changed (i.e. increased or decreased) the expression levels of target genes more than the other types of EPA. This implied that gene expression tended to be activated in monocytes and CD4^+^ T cells, but repressed in H1-hESC and iPSC by enhancers.EPA shortened at forward–reverse orientation of CTCF-binding sites also showed the highest normalized number of functional enrichments of transcriptional target genes, as shown in the previous paragraphs.

## Discussion

Genome-wide functional enrichments and gene expression levels of putative target genes of human transcription factors were investigated. Human putative transcriptional target genes showed significantly larger numbers of functional enrichments than gene expression information alone, independent of transcriptional target genes. Moreover, when the number of functional enrichments of human putative transcriptional target genes was normalized by the total number of transcriptional target genes, native putative transcriptional target genes showed the highest ratio of functional enrichments, compared with target genes partially including randomly selected genes. The ratio of functional enrichments was decreased according to the increase of the ratio of randomly selected genes in target genes. These tendencies were observed in putative transcriptional target genes predicted from both open chromatin regions and ChIP-seq data of transcription factors. Prediction of transcriptional target genes from open chromatin regions includes false positives, since DNase I cleavage bias affects the computational analysis of DNase-seq experiments [14]. However, the detection of ChIP-seq peaks is also changed depending on the methods to identify them and the depth of DNA sequencing of ChIP-seq experiments [15]. Though human putative transcriptional target genes include false positives, they showed significantly the largest number of functional enrichments, compared with target genes including 5 – 60% of randomly selected genes (Figure 2).

The median expression level of human putative transcriptional target genes was changed according to the criteria of enhancer-promoter assignments, and was correlated with the normalized number of functional enrichments. The median expression level of transcriptional target genes was ‘decreased’ significantly in transcriptional target genes predicted using enhancers, compared with those predicted using promoters in H1-hESC and iPSC, and the median expression level was ‘increased’ significantly in target genes predicted using enhancers, compared with those predicted using promoters in monocytes and CD4^+^ T cells. These results implied that transcription factors bound in enhancers act as repressors in H1-hESC (ES) and iPSC, but those act as activators in monocytes and CD4^+^ T cells. The change of functional roles of transcription factors depending on the cell types would be analyzed and reported elsewhere.

The median expression level was increased significantly in target genes predicted using enhancers, compared with those predicted from promoters in immune cells using gene expression data (Blueprint RNA-seq RPKM data; GSE58310), but smaller number of target genes showed the increase of median expression level using gene expression data (ENCODE; GSM984609). The results of the analyses may be slightly different depending on gene expression data. H1-hESC (ES) and iPSC showed a strong tendency of decrease of median expression levels of transcriptional target genes between enhancers and promoters.

The gene symbols of transcription factors were sometimes different among databases, because more than one gene symbol are assigned to some transcription factors and some gene symbols are spelled in several different ways. These differences need to be identified with manual curations. This analysis will be required to predict transcriptional cascades by associating transcription factors with transcriptional target genes consisting of transcription factors. In the analyses of transcriptional cascades, to reduce false positive predictions of enhancer-promoter associations from open chromatin regions, the identification of DNase peaks will be modified using a new tool such as HINT [16].

In this study, I focused on three types of immune cells and stem cells such as H1-hESC and iPSC to examine transcriptional target genes in a genome scale, since in my previous study, I examined transcriptional cascades involved in the differentiation of immune cells as introduced in Background [5]. Furthermore, I confirmed the features of functional enrichments of putative transcriptional target genes are commonly found in other four types of normal and disease cells (HUVEC, IMR90, MCF-7 and HMEF).

It is difficult to predict enhancer-promoter associations using a single parameter, so that machine learning methods to combine several parameters have been proposed [17-19]. These methods showed high accuracy in predicting enhancer-promoter associations (I tried to use some of the tools, but they did not work properly. I am waiting for the authors to update the tools). However, molecular mechanisms of enhancer-promoter interactions are not clearly understood. CTCF has been found to bind at chromatin interaction anchors and form chromatin interactions [12]. About 20-40% of chromatin interaction anchors included DNA binding sequences of CTCF, when I examined public Hi-C experimental data [20, 21]. Among 33,939 RefSeq transcripts, 7,202 (21%), 4,404 (13%), and 6,921 (20%) (*p*-value < 10^-5^ in the search for CTCF-binding motifs using FIMO) to 9,608 (28%), 5,806 (17%), and 9,137 (27%) (*p*-value < 10^-4^) of transcripts had forward–reverse orientation of CTCF-binding sites within 1 Mb from transcriptional start sites in the three immune cell types, respectively. These analyses implied that other factors might be involved in chromatin interactions. ZNF143 has been reported to locate at promoter regions of chromatin interaction anchors [22]. To predict the other factors and molecular mechanisms, the analyses in this study would be useful to examine further the criteria in predicting enhancer-promoter associations. Machine learning methods need the information what parameters should be used for prediction, so it would be better to choose parameters involved in predicting enhancer-promoter associations. To improve the prediction and understand the molecular mechanisms of enhancer-promoter interactions, I am promoting the analyses of chromatin interaction anchors, and the results of the analyses will be reported elsewhere.

## Conclusion

In this study, human transcriptional target genes were predicted using open chromatin regions, ChIP-seq data, and DNA binding sequences of transcription factors in databases. Human putative transcriptional target genes showed significant functional enrichments. Journal papers showed that frequent functional enrichments were related to the cellular functions. The normalized number of functional enrichments was the highest in native putative transcriptional target genes, compared with target genes partially replaced with randomly selected genes. The normalized number of functional enrichments of human putative transcriptional target genes changed according to the criteria of enhancer-promoter assignments and correlated with the median expression level of the target genes. The normalized numbers of functional enrichments of transcriptional target genes did not show the highest number in the criterion of enhancer-promoter assignments covering the longest distance from transcriptional start site among four criteria. This suggested that there is a criterion of enhancer-promoter assignments that shows the highest normalized number of functional enrichments. The median expression level of transcriptional target genes was ‘decreased’ significantly in transcriptional target genes predicted using enhancers, compared with those predicted using promoters in H1-hESC and iPSC, and the median expression level was ‘increased’ significantly in target genes predicted using enhancers, compared with those predicted using promoters in immune cells. These results implied that transcription factors bound in enhancers act as repressors in H1-hESC (ES) and iPSC, but those act as activators in immune cells. These analyses and characters of human putative transcriptional target genes would be useful to examine the criteria of enhancer-promoter assignments and to predict the novel mechanisms and factors such as DNA binding proteins and DNA sequences of enhancer-promoter interactions.

## Methods

### Searches for transcription factor binding sequences from open chromatin regions

To examine transcriptional regulatory target genes, bed files of hg19 narrow peaks of ENCODE DNase-DGF and DNase data for Monocytes-CD14^+^_RO01746 (GSM1024791; UCSC Accession: wgEncodeEH001196), CD4^+^_Naive_Wb11970640 (GSM1014537; UCSC Accession: wgEncodeEH003156), CD20^+^_RO01778 (GSM1014525; UCSC Accession: wgEncodeEH002442), H1-hESC (GSM816632; UCSC Accession: wgEncodeEH000556), iPSC (GSM816642; UCSC Accession: wgEncodeEH001110), HUVEC (GSM1014528; UCSC Accession: wgEncodeEH002460), IMR90 (GSM1008586; UCSC Accession: wgEncodeEH003482), MCF-7 (GSM816627; UCSC Accession: wgEncodeEH000579), and HMEC (GSM816669; UCSC Accession: wgEncodeEH001101) from the ENCODE website (http://hgdownload.cse.ucsc.edu/goldenPath/hg19/encodeDCC/wgEncodeUwDgf/) were used. For comparison with transcriptional target genes predicted using ChIP-seq data, bed files of hg19 narrow peaks of ENCODE ChIP-seq data for 19 transcription factors (TF) (BACH1, BRCA1, C/EBPbeta, CHD2, c-JUN, c-MYC, GTF2I, JUND, MAFK, MAX, MXI1, NRF1, RAD21, RFX5, SIN3A, SUZ12, TBP, USF2, ZNF143) in H1-hESC from the ENCODE website (https://genome.ucsc.edu/cgi-bin/hgFileUi?db=hg19&g=wgEncodeAwgTfbsUniform) were utilized.

To identify transcription factor binding sites (TFBS) from the DNase-DGF data, TRANSFAC (2013.2), JASPAR (2010), UniPROBE, BEEML-PBM, high-throughput SELEX, Human Protein-DNA Interactome, and transcription factor binding sequences of ENCODE ChIP-seq data were used [23-29]. Position weight matrices of transcription factor binding sequences were transformed into TRANSFAC matrices and then into MEME matrices using in-house Perl scripts and transfac2meme in MEME suite [30]. Transcription factor binding sequences of transcription factors derived from vertebrates were used for further analyses. Searches were conducted for transcription factor binding sequences from the central 50-bp regions of each narrow peak using FIMO with *p*-value threshold of 10^-5^ [31]. Transcription factors corresponding to transcription factor binding sequences were searched computationally by comparing their names and gene symbols of HGNC (HUGO Gene Nomenclature Committee) -approved gene nomenclature and 31,848 UCSC known canonical transcripts(http://hgdownload.cse.ucsc.edu/goldenpath/hg19/database/knownCanonical.txt.gz), as transcription factor binding sequences were not linked to transcript IDs such as UCSC, RefSeq, and Ensembl transcripts.

### Prediction of transcriptional target genes

Target genes of a transcription factor were assigned when its TFBS was found in DNase-DGF narrow peaks in promoter or extended regions for enhancer-promoter association of genes (EPA). Promoter and extended regions were defined as follows: promoter regions were those that were within distances of ±5 kb from transcriptional start sites (TSS). Promoter and extended regions were defined as per the following four association rules, which are similar or same as those defined in a previous study [11]: (1) the basal plus extension association rule assigns a basal regulatory domain to each gene regardless of other nearby genes. The domain is then extended to the basal regulatory domain of the nearest upstream and downstream genes, and includes a 5 kb + 5 kb basal region and an extension up to 300 kb or the midpoint between the TSS of the gene and that of the nearest gene upstream and downstream; (2) 5 kb + 1 kb basal region and an extension up to 1 Mb; (3) the two nearest genes association rule, which extends the regulatory domain to the TSS of the nearest upstream and downstream genes without the limitation of extension length; and (4) the single nearest gene association rule, which extends the regulatory domain to the midpoint between the TSS of the gene and that of the nearest gene upstream and downstream without the limitation of extension length. Association rule (1) was used in our previous study [5]. Association rule (2), (3), and (4) were the same as those in Figure 3A of the previous study [11], however, association rules (3) and (4) did not have the limitation of extension length in this study. The genomic positions of genes were identified using ‘knownGene.txt.gz’ file in UCSC bioinformatics sites [32]. The file ‘knownCanonical.txt.gz’ was also utilized for choosing representative transcripts among various alternate forms for assigning promoter and extended regions for enhancer-promoter association of the genes. From the list of transcription factor binding sequences and transcriptional target genes, redundant transcription factor binding sequences were removed by comparing the target genes of a transcription factor binding sequence and its corresponding transcription factor; if identical, one of the transcription factor binding sequences was used. When the number of transcriptional target genes predicted from a transcription factor binding sequence was less than five, the transcription factor binding sequence was omitted.

### Gene expression analyses

For gene expression data, RNA-seq reads mapped onto human hg19 genome sequences were obtained, including ENCODE long RNA-seq reads with poly-A of monocytes CD14^+^ cells, CD20^+^ B cells, H1-hESC, iPSC, HUVEC, IMR90, MCF-7, and HMEC (GSM984609, GSM981256, GSE26284, GSM958733, GSM2344099, GSM2344100, GSM958734, GSM2400222, GSM765388, and GSM758571), and UCSF-UBC human reference epigenome mapping project RNA-seq reads with poly-A of naive CD4^+^ T cells (GSM669617). Two replicates were present for monocytes CD14^+^ cells, CD20^+^ B cells, H1-hESC, iPSC, HUVEC, IMR90, MCF-7, and HMEC and a single one for CD4^+^ T cells. RPKMs of the RNA-seq data were calculated using RSeQC [33]. For monocytes, Blueprint RNA-seq RPKM data (GSE58310, GSE58310_GeneExpression.csv.gz,Monocytes_Day0_RPMI) was also used [34]. Based on RPKM, UCSC transcripts with expression levels among top 30% of all the transcripts were selected in each cell type.

### Functional enrichment analyses

The functional enrichments of target genes of a TFBS and its corresponding transcription factor were examined using GO-Elite v1.2.5 with *p*-value threshold at 1, and after GO-Elite analyses a false discovery rate (FDR) test was performed with *q*-value threshold at 10^-3^ to correct for multiple comparisons of thousands of groups of transcriptional target genes in each cell type and condition [35]. For examining functional enrichments of high or low expressed genes independent of transcriptional target genes, the *p*-value threshold was set to 0.01 or 0.05 to confirm that the results were not significantly changed. UCSC gene IDs were transformed into RefSeq IDs prior to GO-Elite analyses. GO-Elite uses 10 databases for identifying functional enrichments: (1) Gene Ontology, (2) Disease Ontology, (3) Pathway Commons, (4) GO Slim, (5) WikiPathways, (6) KEGG, (7) Transcription factor to target genes, (8) microRNA to target genes, (9) InterPro and UniProt functional regions (Domains), and (10) Cellular biomarkers (BioMarkers). To calculate the normalized numbers of functional enrichments of target genes, the numbers of functional enrichments were divided by the total number of target genes in each cell type and condition, and were multiplied by 10^5^. In tables showing the numbers of functional enrichments in 10 databases, heat maps were plotted according to *Z*-scores calculated from the numbers of functional enrichments of each database using in-house Excel VBA scripts. In the comparisons of the normalized numbers of functional enrichments of target genes in cell types and conditions, if the number of a functional annotation in a cell type or condition was two times larger than that in the other cell type or condition, the functional annotation was recognized as more enriched than the other cell type or condition.

To investigate whether the normalized numbers of functional enrichments of transcriptional target genes correlate with the prediction of target genes, a part of target genes were changed with randomly selected genes with high expression level (top 30% expression level), and functional enrichments of the target genes were examined. First, 5%, 10%, 20%, 40%, and 60% of target genes were changed with randomly selected genes with high expression level in monocytes, CD4^+^ T cells, and CD20^+^ B cells. Second, as another randomization of target genes, the same number of 5%, 10%, 20%, 40%, and 60% of target genes were selected randomly from highly expressed genes, then added them to the original target genes, and functional enrichments of the target genes were examined. All analyses were repeated three times to estimate standard errors (Figure 2A and B, Figure S1 and S2, and Table S1). The same analysis was performed using DNase-DGF data and ChIP-seq data of 19 TF in H1-hESC. Transcriptional target genes were predicted from promoter (Tables S2).

### CTCF-binding sites

CTCF ChIP-seq data for monocytes CD14^+^ cells (GSM1003508_hg19_wgEncodeBroadHistoneMonocd14ro1746CtcfPk.broadPeak.gz), CD4^+^ T cells (SRR001460.bam), CD20^+^ B cells (GSM1003474_hg19_wgEncodeBroadHistoneCd20CtcfPk.broadPeak.gz), H1-hESC (wgEncodeAwgTfbsUtaH1hescCtcfUniPk.narrowPeak.gz), iPSC (GSE96477), HUVEC (wgEncodeAwgTfbsUwHuvecCtcfUniPk.narrowPeak.gz), IMR90 (wgEncodeAwgTfbsSydhImr90CtcfbIggrabUniPktfbsf.narrowPeak.gz), MCF-7 (wgEncodeAwgTfbsUwMcf7CtcfUniPktfbsf.narrowPeak.gz), and HMEC (wgEncodeAwgTfbsUwHmecCtcfUniPktfbsf.narrowPeak.gz) were used.SRR001460.bam was sorted and indexed by SAMtools and transformed into a bed file using bamToBed of BEDTools [36, 37]. ChIP-seq peaks were predicted by SICER-rb.sh of SICER with optional parameters ‘hg19 1 200 150 0.74 200 100’ [38]. Extended regions for enhancer-promoter association (association rule 4) were shortened at the genomic locations of CTCF-binding sites that were the closest to a transcriptional start site, and transcriptional target genes were predicted from the shortened enhancer regions using TFBS. Furthermore, promoter and extended regions for enhancer-promoter association (association rule 4) were shortened at the genomic locations of forward–reverse orientation of CTCF-binding sites. When forward or reverse orientation of CTCF-binding sites were continuously located in genome sequences several times, the most external forward–reverse orientation of CTCF-binding sites were selected.

## Abbreviations

TF: transcription factors
TFBS: transcription factor binding sites
TSS: transcriptional start sites
ENCODE: encyclopedia of DNA elements
ChIP-seq: ChIP-sequencing, chromatin immunoprecipitation followed by massively parallel DNA sequencing
RNA-seq: RNA-sequencing
EPA: promoter and extended regions for enhancer-promoter association
FR: forward-reverse
RF: reverse-forward
FF: forward-forward
RR: reverse-reverse

## Declarations

### Ethics approval and consent to participate

I used experimental data in public databases, so this is not applicable.

### Consent for publication

Not applicable

### Availability of data and material

I used experimental data in public and commercial databases, so this is not applicable. TRANSFAC, which is a database of transcription factors, is commercial database.

### Authors’ contributions

NO designed and performed the research

## Acknowledgements

The supercomputing resource was provided by Human Genome Center of the Institute of Medical Science at the University of Tokyo. Computations were partially performed on the NIG supercomputer at ROIS National Institute of Genetics.

## Authors’ information

### References used in Table 4

Monocytes: IFN-gamma pathway[39], IL3-mediated signaling events[40], GMCSF-mediated signaling events[41], PDGF receptor signaling network[42], VEGF and VEGFR signaling network[43]

CD4+ T cell: Integrin family cell surface interactions[44], IFN-gamma pathway[45], LKB1 signaling events[46], TCR signaling in naive CD4+ T cells[47], IL3-mediated signaling events[48]

CD20+ B cell: Integrin family cell surface interactions[49], IL5-mediated signaling events[50], Insulin Pathway[51], mTOR signaling pathway[52], Sphingosine 1-phosphate (S1P) pathway[53]

### References used in Table 6

Monocytes: CXCR4-mediated signaling events[54], GPVI-mediated activation cascade[55], Signaling events mediated by TCPTP[56], p38 MAPK signaling pathway[57], IL4-mediated signaling events[58]

CD4+ T cell: CXCR4-mediated signaling events[59], Fc-epsilon receptor I signaling in mast cells[60], JNK signaling in the CD4+ TCR pathway[61], Reelin signaling pathway[62], Semaphorin interactions[63]

CD20+ B cell: Syndecan-4-mediated signaling events[64], CXCR4-mediated signaling events[65], EPO signaling pathway[66], Stabilization and expansion of the E-cadherin adherens junction[67], Interleukin-1 signaling[68]

## Endnotes

## Tables

Tables are attached in separate PDF files (Table 1 – 6) in Attachment folder, because the tables are colored to understand them easily and statistically different numbers are shown with lines and asterisks.

